# Elemental diet enriched in amino acids alters gut microbial community and prevents colonic mucus degradation in mice with colitis

**DOI:** 10.1101/2022.06.01.494459

**Authors:** Bowei Zhang, Congying Zhao, Yunhui Zhang, Xuejiao Zhang, Xiang Li, Xiaoxia Liu, Jia Yin, Xinyang Li, Jin Wang, Shuo Wang

## Abstract

The role of dietary amino acids or intact proteins in the progression of colitis remains controversial, and the mechanism involvin<u>g</u> gut microbes is unclear. Here, we investigated the effects of an elemental diet (ED) enriched in amino acids and a polymeric diet enriched in intact protein on the pathogenesis of DSS-induced colitis in mice. Our results showed that ED induced remission of colitis in mice. Notably, ED treatment reduced the abundance of the mucolytic bacteria *Akkermansia* and *Bacteroides*, which was attributed to the decreased colonic protein fermentation. Consistently, the activities of mucolytic enzymes were decreased, leading to the protection of mucus layer degradation and microbial invasion. The fecal microbiota transplantation of ED-fed mice reshaped microbial ecology and alleviated intestinal inflammation in recipient mice. ED failed to induce remission of colitis in pseudo-germ-free mice. Together, we convincingly demonstrated the critical role of gut microbiota in the prevention of ED on colitis.

**Importance:** The prevalence of inflammatory bowel disease is rapidly increasing and has become a global burden. Several specific amino acids have been shown to benefit mucosal healing and colitis remission. However, the role of amino acids or intact proteins in diets and enteral nutrition formulas is controversial, and the mechanisms involving gut microbes remain unclear. In this study, we investigated the effects of an elemental diet (ED) enriched in amino acids and a polymeric diet enriched in intact protein on the pathogenesis of colitis in mice. The underlying mechanisms were explored by utilizing fecal microbiota transplantation and pseudo-germ-free mice. ED treatment reduced the abundance of mucolytic bacteria, thereby protecting the mucus layer from microbial invasion and degradation. For the first time, we convincingly demonstrated the critical role of gut microbiota in the effects of ED. This study may provide new insights into the gut microbiota-diet interaction in human health.

## Introduction

Inflammatory bowel disease (IBD), including ulcerative colitis (UC) and Crohn’s disease (CD), is characterized by abdominal pain, diarrhea, pus, and blood in the stool (1). The prevalence of IBD is highest in Europe and North America, with a rapidly increasing trend in developing countries. The incidence of the disease gradually tends to be younger (2), and has become a global burden (3).

The pathological changes of IBD mainly occur in the colonic mucosa and submucosa, and gradually spread to the entire colon (4). The disorder of intestinal flora and the damage of intestinal mucosa is the main features of IBD, and are important factors for the aggravation of intestinal inflammation (5, 6). The mucus layer of the colon and intestine is the first barrier that protects the gut from bacteria (7). The colonic mucus layer is divided into an outer mucus layer and an inner mucus layer, the trunk of which is mainly composed of MUC2 secreted by goblet cells (8, 9). A key nutritional feature of the intestinal mucus layer is the high content of polysaccharide, of which the content of O-glycan is up to 80% (10). However, only a subset of gut microbes are capable of utilizing this nutrient source (11). In a healthy state, bacteria adhere to the outer mucus layer. Some bacteria such as *Akkermansia muciniphila* (*A.muciniphila*) and *Bacteroides*, can degrade mucus and produce short-chain fatty acids, which could provide energy for goblet cells and promote mucus secretion to maintain the sterile state of the inner mucus layer (12, 13). However, in the case of colitis, goblet cells are destroyed and mucolytic bacteria proliferate. This results in a thinning of the intestinal mucus layer, allowing bacteria to invade the intestinal epithelium, leading to a more severe inflammatory response (6, 14). Currently, several treatments for colitis are available, including non-targeted therapies such as 5-aminosalicylic acid (5-ASA), glucocorticoids and immunosuppressants (azathioprine) (15), as well as targeted biologics including anti-tumor necrosis factor (TNF) therapy and c-Jun N-terminal kinase (JNK) inhibitors (16). However, long-term use of these drugs may cause side effects. The pathogenesis of IBD involves both genetic and environmental factors. Among that, diet plays an important role in the progression of IBD (17). Dietary intervention strategies are more acceptable to patients and have fewer side effects, and it has attracted widespread attention (18).

Nutrients act as critical regulators of the immune system and gut microbial ecology (19). Among the nutrients, several specific dietary amino acids participate in cellular and microbial metabolic pathways and play a role in mucosal healing and gut microbiota shaping (20, 21). An amino acid-enriched diet reduces dietary antigens in the gut lumen and is generally considered to be better absorbed in the proximal small intestine, while the residual amount in the distal small intestine and colon is minimal (22). Previous studies have fed colitis mice with an amino acid-based elemental diet (ED) and a polymeric diet with intact proteins, respectively, and found that the amino acid diet inhibited colon inflammation in the mice and suppressed Th1 and Th17 cell responses (23, 24). Moreover, as a formula for enteral nutrition (EN), ED has been shown to induce remission of IBD in patients (18). At the same time, studies comparing different EN treatments have yielded conflicting results. Earlier researches suggest that CD patients treated with ED have significantly higher remission rates than those on a polymeric diet (25). But later studies found that ED was equally effective as polymeric diets (26). Given the critical role of gut microbes in the progression of IBD, these inconsistent results may involve individual differences in gut microbes. ED can alter the gut microbial community by altering the nutrient composition of the gut microbiota (18), but the underlying mechanisms by which ED alleviates IBD is still lacking.

Therefore, this study aims to investigate the effect of ED on the progression of colitis in mice. By utilizing fecal microbiota transplantation and antibiotic-treated pseudo-sterile mouse models, we elucidated the molecular mechanisms by which the ED diet alters gut microbiomes and affects the progression of colitis.

## Results

### Elemental diet prevented the progression of colitis in mice

To evaluate the effect of ED on colitis, mice were fed with ED for two weeks and then given DSS for 3 cycles (Fig. 1A). During the final stage of the experiment, the disease activity index (DAI) of the DSS-treated mice gradually increased. The mice developed severe diarrhea and blood in the stool, and their body weight was significantly reduced (Fig. 1B and 1C). The intervention of ED significantly alleviated the above symptoms. The shortened colon length is an important indicator of colitis. DSS significantly reduced the colon length in mice but was effectively prevented by ED (Fig. 1D and 1E). Histopathology showed that colitis mice showed crypt deformation, epithelial damage, and obvious infiltration of inflammatory cells in the submucosa, which were improved by ED treatment (Fig. 1F and 1G). DSS-induced colitis significantly increased the neutrophil marker myeloperoxidase (MPO) but decreased significantly by ED (Fig. 1H). These data suggested that ED intervention can effectively improve the pathological damage of DSS-induced colitis. Interestingly, there was no significant difference in colitis-related parameters between the two diets in normal mice, but the body weight of the AA group mice was slightly increased (Fig. 1B).

**Figure 1.**
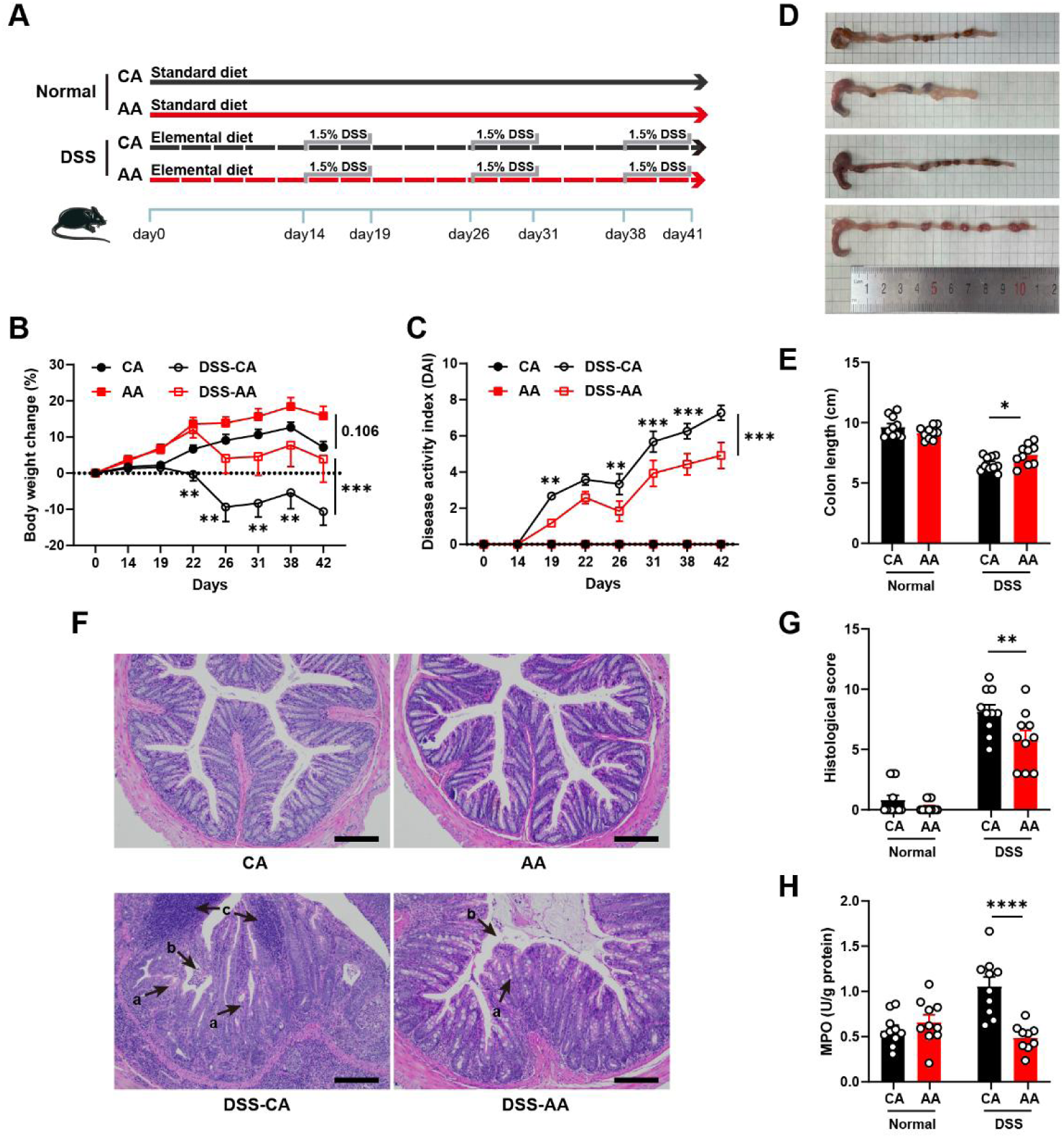
Elemental diet prevents the progression of DSS-induced colitis in mice. Schematic illustration of the experimental design (A). Body weight (B), disease activity index (C), colon photos (D), colon length (E). HE-stained sections of colon tissue, arrows a showed crypt deformation, arrows b showed epithelial damage, arrows c showed infiltration of inflammatory cells in the submucosa, 100× magnification, bar = 200 μm (F). Histological score (G) and colonic MPO activity (H). Data were represented as mean ± SEM, and two-way ANOVA followed by Bonferroni multiple comparison test (n=10), was used to test significance \**p* < 0.05, \*\**p* < 0.01, or \*\*\**p* < 0.001.

### Elemental diet inhibited intestinal inflammation in mice

We then examined the mRNA expression of colitis-related proteins to explore the mechanism by which ED alleviates colitis. The results showed that the expressions of pro-inflammatory factors, including IL-6, TNF-α, IFN-γ, IL-12, and IL-23 were significantly down-regulated in the DSS-AA group compared with the DSS-CA group (Fig. 2A). The expression of IL-1β, which is associated with inflammasome activation, was significantly decreased in the DSS-AA-treated group, but the expression of IL-17, which is associated with Th17 cell differentiation, was not significantly altered.

**Figure 2.**
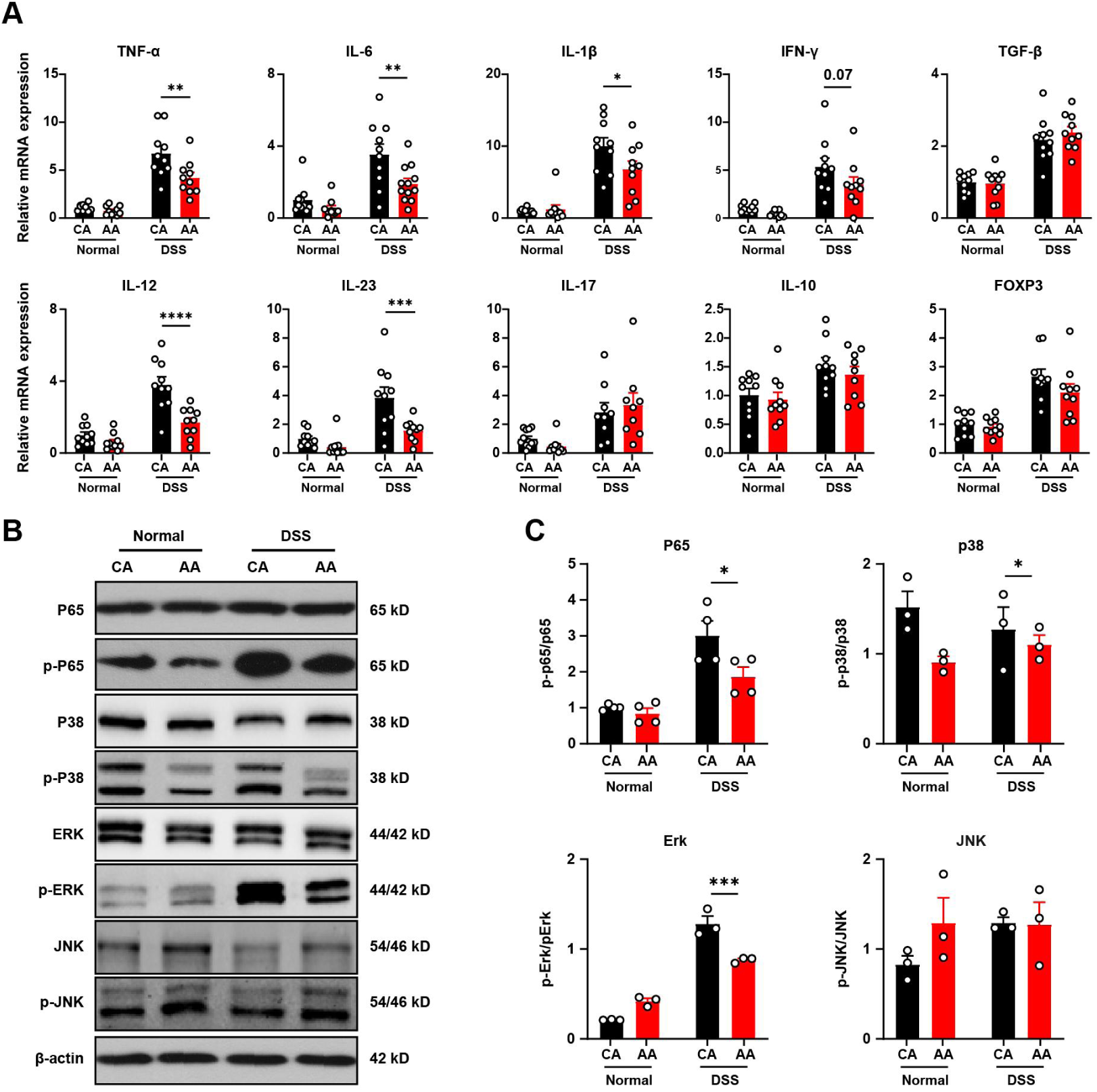
Elemental diet inhibits intestinal inflammation in mice. Relative mRNA expressions of cytokines in colon tissue (A). Expression of NFκB and MAPK pathway proteins in colon tissues (B and C). Data were represented as mean ± SEM, and two-way ANOVA followed by Bonferroni multiple comparison test (n=10 for A, and n=3-4 for B and C), \**p* < 0.05, \*\**p* < 0.01, or \*\*\**p* < 0.001.

Western blot results showed that the phosphorylation levels of p65, Erk, and p38 in the DSS-AA group were significantly lower than those in the DSS-CA group (Fig. 2B and 2C). This indicates that ED could inhibit the activation of inflammatory signaling pathways NF-κB and MAPK, thereby reducing the release of downstream inflammatory factors and improving colitis in mice.

### Elemental diet increased mucin expression

Disruption of the epithelial barrier is a key driver of intestinal inflammation. We observed that the serum level of LPS in the DSS-AA group was decreased compared with the DSS-CA group (Fig. 3A). Therefore, we further examined the mRNA expression of intestinal barrier-related proteins. In the DSS-AA group, the mRNA expressions of antibacterial peptide RegIIIγ secreted by Paneth cells were significantly increased (Fig. 3B). However, the mRNA expression of most key proteins such as MUC2, occludin, ZO-1, and claudin was comparable between the two groups.

**Figure 3.**
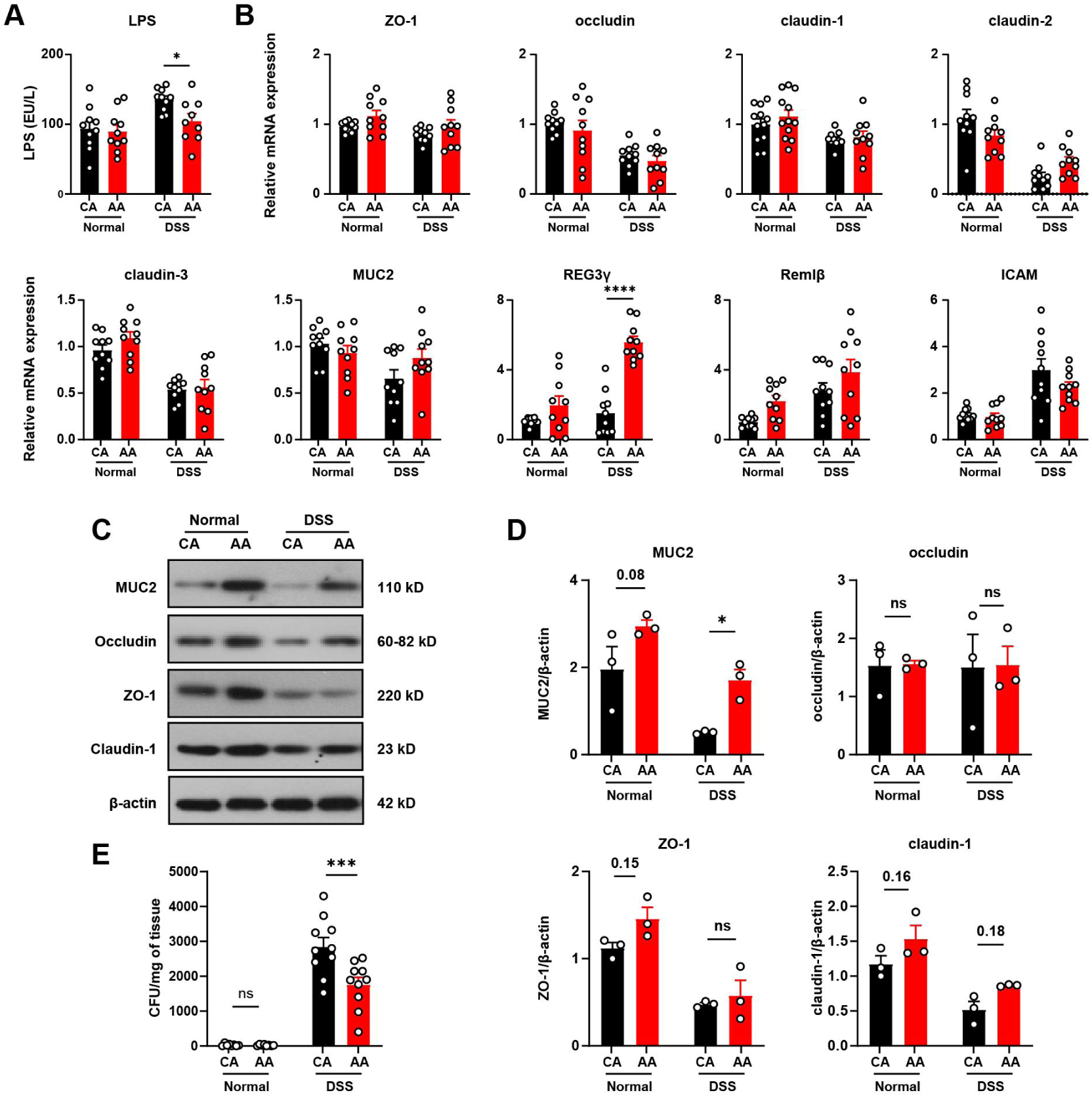
Elemental diet increases mucin expression. LPS level in serum (A). Relative mRNA expression of gut barrier-related proteins (B). The expression level of gut barrier-related protein (C and D). Quantification of aerobic bacteria in mesenteric lymph node (E). Data were represented as mean ± SEM, and and two-way ANOVA followed by Bonferroni multiple comparison test (n=10 for A, B and E, n=3 for C and D), \**p* < 0.05, \*\**p* < 0.01, or \*\*\**p* < 0.001.

According to the results of Western blot, there was no significant difference in the expression of ZO-1, occludin, and Claudin-1 between the DSS-AA group and the DSS-CA group. Interestingly, the amino acid-based ED increased the protein expression of MUC2 in both normal and colitis mice (Fig. 3C and 3D). MUC2 is a major component of the colonic mucus layer, which helps to resist the invasion of intestinal pathogens. Correspondingly, we observed decreased microbial translocation in the mesenteric lymph node of ED-fed mice (Fig. 3E). The above results indicated that the improvement of colitis by ED was mediated by the regulation of intestinal mucins, and this did not involve changes at the transcriptional level.

### Elemental diet reduced mucolytic bacteria abundance and mucolytic enzyme activity

To further explore the underlying mechanism, we determined the relative abundance of gut microbiota by meta-genomic sequencing. According to the results of principal coordinate analysis (PCoA), the samples in the DSS-CA group were distinguished from those in the DSS-AA group (Fig. 4 and Fig S1). This difference was also observed between the CA Group and the AA group, suggesting that ED altered the composition of gut microbiota.

**Figure 4.**
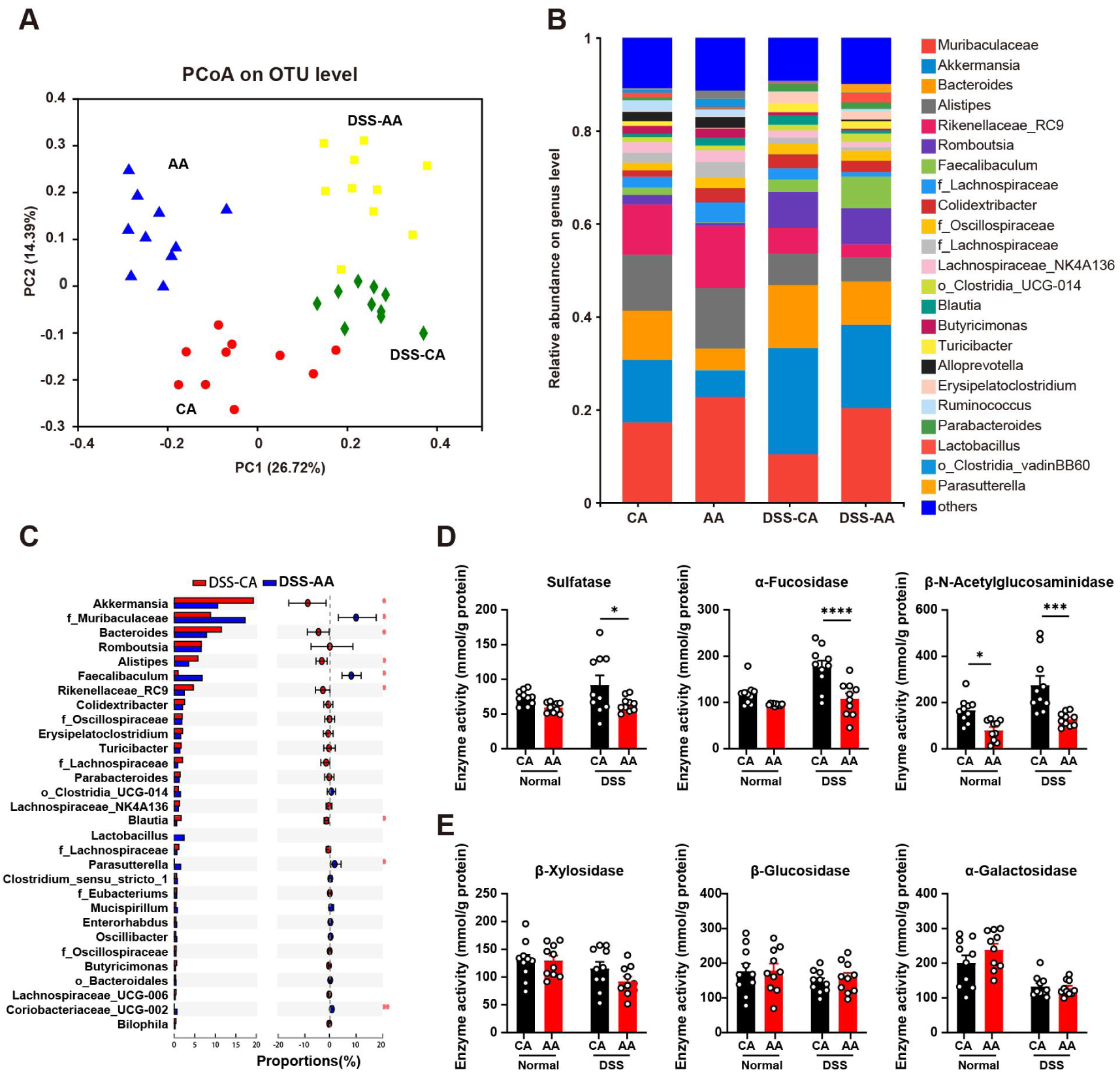
Elemental diet reduced mucolytic bacteria abundance and mucolytic enzyme activity. Principal coordinate analysis (PCoA) of fecal microbiota at OTU level (A). Relative abundance of gut microbes at the genus level (B). Difference between gut microbiota in the DSS-CA group and the DSS-AA group at the genus level (C). Activities of mucolytic enzymes and carbohydrate-active enzymes (D). Data were represented as mean ± SEM, significance was examined by Mann-Whitney test for C (n=10), and was examined using two-way ANOVA followed by Bonferroni multiple comparison tests for D and E (n=10), \**p* < 0.05, \*\**p* < 0.01, or \*\*\**p* < 0.001.

ED decreased the abundance of *Akkermansia* and *Bacteroides*, and increased the abundance of Muribaculaceae and *Faecalibaculum* in both the DSS-treated and normal groups (Fig. 4B, 4C, and Fig. S2). The abundance of these microbes in the samples was 13.7%, 9.1%, 18.4%, and 2.5%, respectively. Furthermore, ED decreased the abundance of *Alistipes*, *Rikenellaceae RC9*, and *Blautia*, and increased the abundance of *Parasutterella* and *Coriobacteriaceae_UCG-002* in DSS-treated mice. The above results showed that ED altered the gut microbial community of mice. Among the microbes with high abundance, *Bacteroide*s and *Akkermansia* were reported to be major contributors to mucus degradation (6).

MUC2 is an O-glycan protein linked by O-glycosidic and disulfide bonds, which can be degraded by microbial mucolytic enzymes and serve as a carbon source for microorganisms. Therefore, we measured the activity of these glycosidases in feces. In the DSS-treated mice, the activities of mucin degrading enzymes, including sulfatase, β-N-acetylglucosaminidase, and α-fucosidase were significantly increased (P <0.05), which was significantly reversed by ED (Fig. 4D). Moreover, the activities of α-galactosidase, β-xylosidase, and β-glucosidase, which are involved in the degradation of plant carbohydrates, were comparable between the CA and the AA groups (Fig. 4E).

In brief, these results suggest that changes in the microbial community, especially mucus-degrading bacteria, may play an important role in the prevention of colitis by ED.

### Prevention of elemental diet on colitis is microbiota dependent

To confirm that gut microbiota plays an important role in the anti-inflammatory effects of ED, we performed fecal microbiota transplantation (FMT) experiments. Donor mice were fed ED or casein diets for two weeks. Recipient mice were treated with antibiotic cocktails for 10 days before gavage to deplete gut microorganisms (Fig. S3). After 14 days of fecal bacteria transplantation, mice were treated with DSS for 7 days to induce colitis (Fig. 5A).

**Figure 5.**
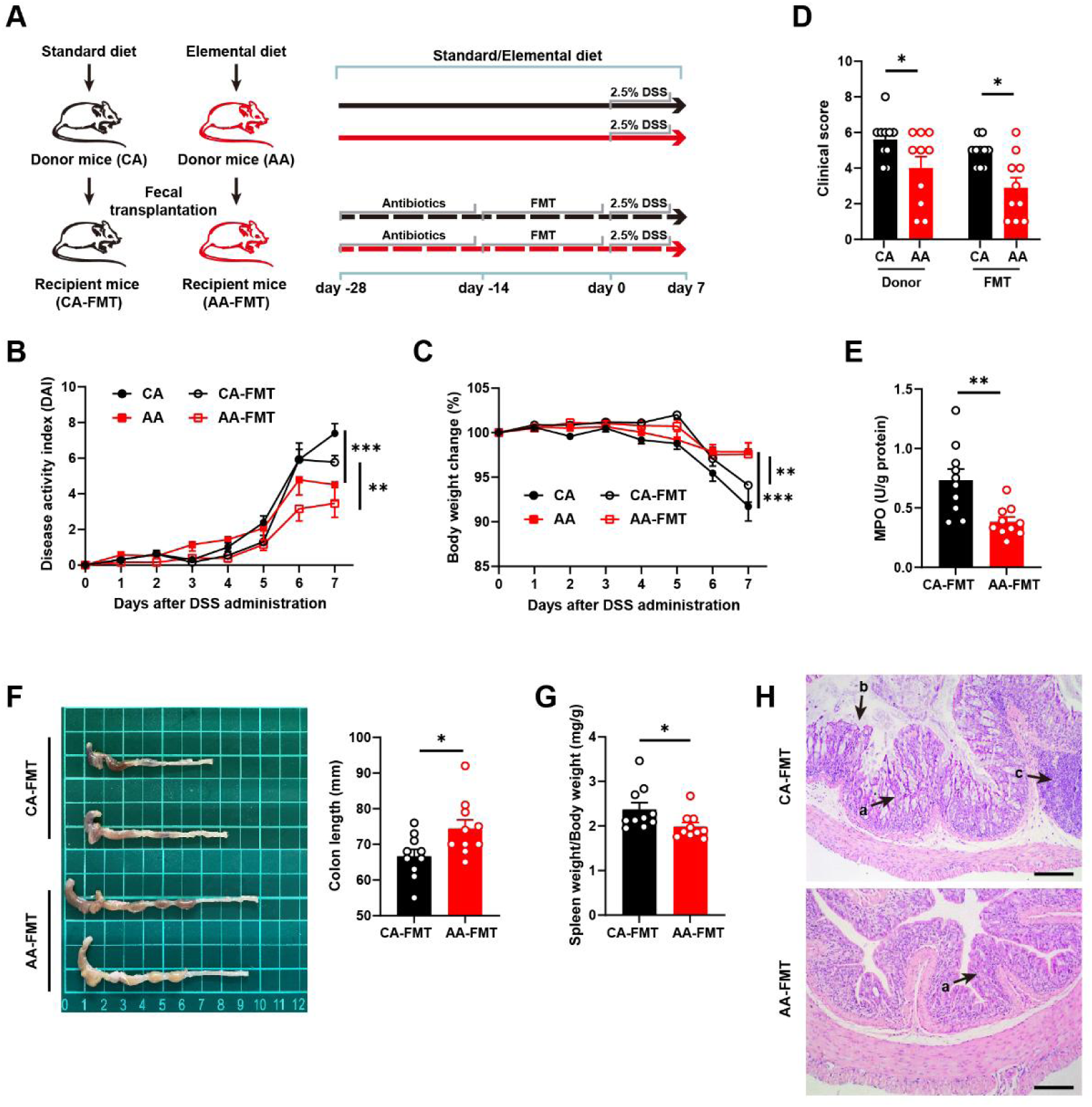
Prevention of elemental diet on colitis is microbiota dependent. Schematic illustration of the experimental design (A). Body weight (B), disease activity index (C), clinical score (D), MPO activity in colon tissue (E). colon length (F), spleen weight (G), and HE staining sections of colon tissue, arrows a showed crypt deformation, arrows b showed epithelial damage, and arrows c showed infiltration of inflammatory cells in the submucosa (H). Data were represented as mean ± SEM, and significance was examined by two-way ANOVA followed by Bonferroni multiple comparison tests for B, C, D (n=10), and was examined by Student’s t-test for E, F, G (n=10). \**p* < 0.05, \*\**p* < 0.01, or \*\*\**p* < 0.001.

Compared with the CA mice, the AA mice had decreased DAI levels and increased colon length (Fig. 5B and 5D), suggesting that ED ameliorated acute colitis in mice caused by a 7-day DSS induction. Compared with the casein receptor group (CA-FMT), the ED receptor group (AA-FMT) had lower DAI and longer colon, along with increased body weight, improved histopathology, and decreased MPO activity (Fig. 5B-H).

In addition, the mRNA expression of inflammatory cytokines Il-6, TNF-α, INF-γ, and IL-1β were decreased in the AA group (Fig. 6A). Western blot results showed that the phosphorylation level of NF-κB p65 in the AA-FMT group was significantly down-regulated compared with the CA-FMT group, and the phosphorylation levels of Erk and p38 in the MAPK pathway were significantly reduced (Fig. 6B and 6C).

**Figure 6.**
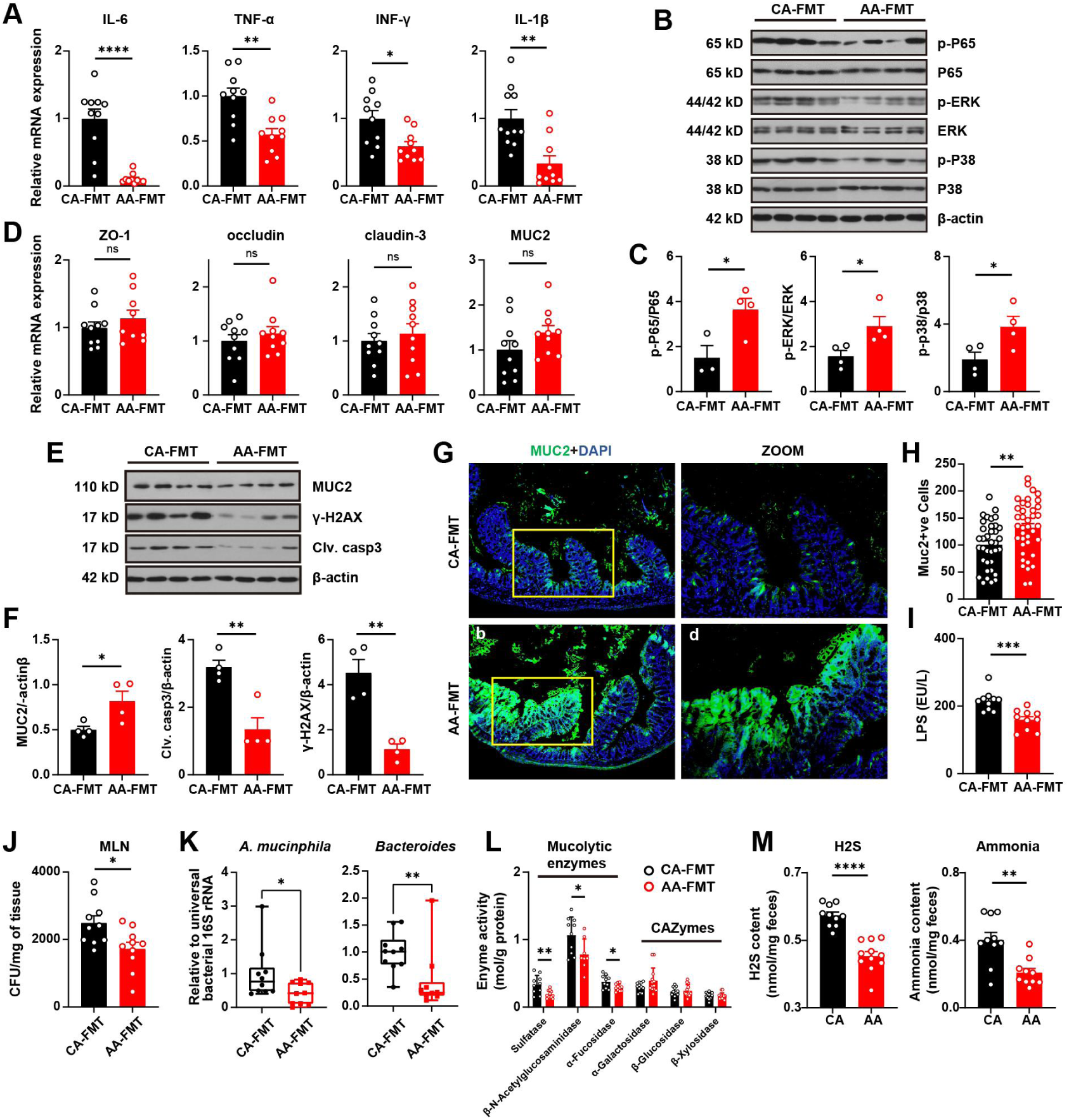
Fecal transplantation of elemental diet-shaped microbiota alleviates erosion of colonic mucus layer. Relative mRNA expression of cytokines in colon tissue (A). The expression level of NFκB and MAPK pathway-related proteins (B and C). Relative mRNA expression of gut barrier-related proteins (D). The expression level of MUC2, gamma H2A histone family member X, and cleaved caspase-3 in colon tissue (E and F). Immunofluorescence analysis of colon sections (G). MUC2-positive cells were counted under a microscope (H). LPS level in serum (I). Quantification of bacterial translocation in the mesenteric lymph node (J). Relative abundance of *A. mucinphila* and *Bacteroides* measured by RT-qPCR (K). Activities of mucolytic enzymes and carbohydrate-active enzymes (L). Fecal content of H_2_S and ammonia (M). Data were represented as mean ± SEM. The significance was examined by Student’s t-test for A, C, D, F, H, I, J, L, and M (n=10), and Mann-Whitney test for K (n=10), \**p* < 0.05, \*\**p* < 0.01, or \*\*\**p* < 0.001.

### Fecal transplantation of elemental diet-shaped microbiota alleviated erosion of the colonic mucus layer

Furthermore, we investigated the effect of fecal bacterial transplantation on the expression of intestinal barrier-related proteins in mice. The results showed that there were comparable differences in the protein and mRNA expression levels of occludin, ZO-1, and claudin-3 between the CA-FMT and the AA-FMT group (Fig. 6D). However, significant upregulation of MUC2 was observed in the AA-FMT group at the protein level (Fig. 6E and 6F). This result was further confirmed by immunofluorescence (Fig. 6G). The number of MUC2-positive goblet cells was significantly increased in the AA-FMT group compared with the CA-FMT group in the immunofluorescence sections (Fig. 6H).

The level of serum LPS was also decreased in the AA-FMT group (Fig. 6I). Correspondingly, bacterial translocation into the mesenteric lymph node (MLN) was significantly inhibited in the AA-FMT group (Fig. 6J). Compared with the CA-FMT group, the expression levels of Clv. caspase-3, the executioner of apoptosis, and DNA damage marker γ-H2AX in colon tissues of the AA-FMT group were significantly reduced (Fig. 6E and 6F).

We then measured the abundance of major mucin-degrading bacteria in the feces of recipient mice. According to the results of RT-qPCR, the relative abundance of *A. mucinphila* and *Bacteroides* was significantly decreased in the FMT-AA group (Fig. 6K). Correspondingly, the activities of mucolytic enzymes were decreased in the FMT-AA group (Fig. 6L), but the activities of CAZymes were not changed.

The elemental formula is thought to reduce food residues reaching the colon (22). To explore the underlying mechanism by which ED affects the microbial community, we measured the content of protein degradation products in feces (Fig. 6M). Notably, donor mice fed the AA diet had significantly lower levels of H_2_S and ammonia in the feces compared to the mice fed the CA diet.

Taking these observations into account, the mechanism by which ED prevents colitis includes reducing the number of mucus-degrading bacteria, thereby inhibiting mucus degradation and suppressing apoptosis and inflammation caused by bacterial translocation. The reduction of proteins entering the large intestine for fermentation plays a key role in the alteration of gut microbiota.

### Prevention of elemental diet against colitis is ineffective in pseudo-germ-free mice

In the end, we compared the effects of ED (ABX-AA) and casein diet (ABX-CA) on colitis in pseudo-germ-free mice (Fig. 7A). Mice were given a broad-spectrum antibiotic cocktail. Bacterial load and fecal DNA concentrations were determined to ensure that gut microbiota was depleted before DSS induction.

**Figure 7.**
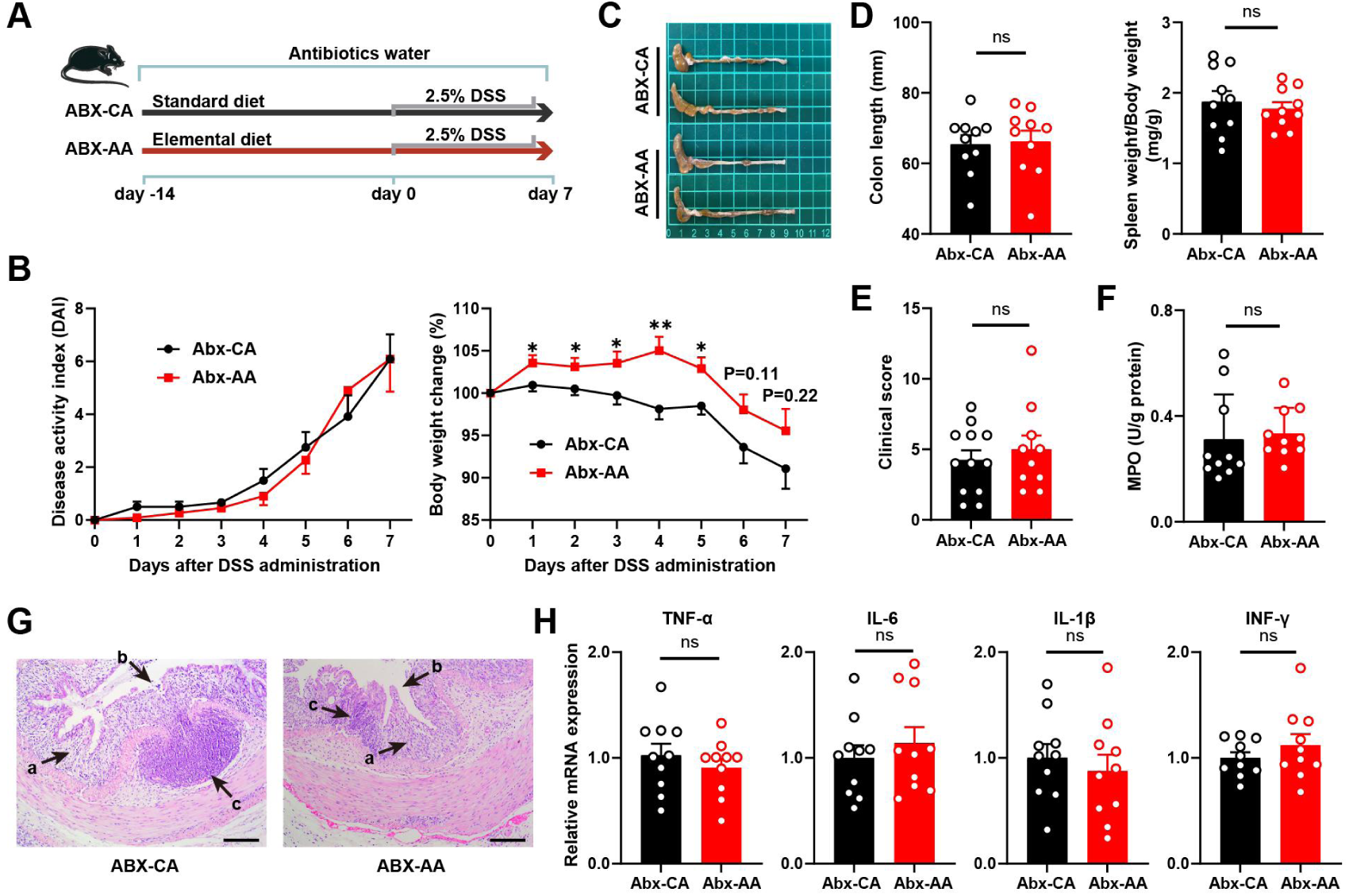
Prevention of elemental diet against colitis is ineffective in antibiotics-treated pseudo-germ-free mice. Schematic illustration of the experimental design (A). Body weight and disease activity index (B), colon photos (C), colon length and spleen weight (D), and clinical score (E). MPO activity in colon tissue (F), HE-stained sections of colon tissue, arrows a showed crypt deformation, arrows b showed epithelial damage, and arrows c showed infiltration of inflammatory cells in the submucosa (G). Relative mRNA expression of cytokines (H). Data were represented as mean ± SEM, and significance was examined by Student’s t-test (n=10), \**p* < 0.05, \*\**p* < 0.01, or \*\*\**p* < 0.001

During the DSS induction period, the body weight was increased in the ABX-AA mice compared with the ABX-CA mice (Fig. 7C). However, the values of DAI and clinical score were comparable between the two groups (Fig. 7B and 7E). In terms of morphology and histopathology, the colon length, edema, crypt morphology, epithelial damage, and inflammatory cell infiltration in the submucosa were comparable in the two groups (Fig. 7C, 7D, and 7G). There was no difference in the activity of inflammation marker MPO (Fig. 7F) in the colon tissue. Similarly, no difference was observed in the expression of pro-inflammatory cytokines IL-1β, IL-6, TNF-α, IFN-γ and the mRNA expression of ZO-1, occludin, Claudin, and MUC2 (Fig. 7H and Fig. S4). The above results indicate that ED has no anti-inflammatory effect in mice deficient gut microbiota.

## Discussion

IBD is a chronic and relapsing inflammatory disease with complex pathogenesis. It is well known that the disorder of gut microbiota is an important factor in the progression of IBD (27). A diet that forms a healthy gut microbial community is an efficient way to modulate host immune response (28), which provides a new strategy for the dietary treatment of IBD. ED therapy has been used as a primary treatment for IBD patients in many parts of the world (18). However, the efficacy of elemental and polymeric formulations remains controversial. The mechanism of ED therapy might be related to alterations in the physical form of the diet, digestion and absorption, and gut microbiota, but the underlying mechanism is not clearly identified. In this study, we compared the effect of an amino acid-enriched ED and an intact protein-enriched polymeric diet on the progression of colitis in mice. We observed the ED-treated mice with increased body weight, decreased DAI, increased colon length, and reduced colon lesions. In conclusion, ED was more effective not only in preventing the progression of chronic colitis but also in preventing the development of acute intestinal inflammation.

Mucus is the first line of defense for intestinal self-protection, physically isolating harmful bacteria (29). Intestinal mucus degradation can lead to colitis in mice (30). MUC2 is the main component of the colonic mucus layer, which forms the skeleton of the mucus (9). It was found that MUC2 knockout mice could cause inflammation that eventually leads to colorectal cancer (31). Therefore, MUC2 is considered a potential therapeutic target for IBD (32). Notably, more than 80% of the mass in mucin is O-glycans, mainly linked by O-glycosidic bonds, and the C- and N- termini are linked by disulfide bonds (9). Some intestinal bacteria utilize mucin as their main carbon source. They can secrete glycosidase and sulfatase to degrade mucin (33). In this study, we found that the degradation of colonic mucus was effectively inhibited in mice fed with ED. At the transcription level, there was no significant difference in MUC2 between the ED group and the casein group, and there was no significant difference in the expression of barrier proteins occludin, claudin, and ZO-1. However, at the protein level, the content of MUC2 in the ED group was significantly increased. Interestingly, this change was eliminated in pseudo-germ-free mice. Meanwhile, the activities of mucus degrading enzymes in feces were detected. It was observed that the activities of mucolytic enzymes were significantly decreased by ED treatment. Consequently, the change in the content of MUC2 is a key mechanism by which ED improves IBD. Reduced mucin-degrading capacity in ED-treated mice is the driving factor affecting MUC2 content, and this change is mediated by intestinal mucolytic bacteria.

The gut microbiota is an important mediator between environmental factors and host health. Short-term changes in diet have been shown to rapidly alter the human gut microbiota (34). The imbalance of gut microbiomes is the main pathogenesis of IBD. The results of our microbiota analysis revealed the reason why ED inhibited the degradation of colonic mucin in mice. *Akkermansia* has received increasing attention in recent years, but its health effects remain controversial. Some studies have found that *Akkermansia* fecal transplantation can extend lifespan in progeria mice (35), and alleviate metabolic diseases such as obesity and diabetes in mice (36). But it is worth noting that numerous studies have shown its negative effects when the host is in a pathological state. For example, the increase of *Akkermansia* in the colon following vitamin deficiency enhances mucolysis, which will lead to intestinal barrier dysfunction and enhance pathogen susceptibility (6). Similarly, *Akkermansia* is found to be abundant in mice with high levels of intestinal inflammation (28), and acts as a pathogen to promote colitis in IL10 (-/-) mice (37). More importantly, clinical research has revealed that IBD patients have higher levels of mucolytic bacteria (38), of which *Akkermansia* and *Bacteroides* are the main members (6). In this study, the relative abundance of *Akkermansia* and *Bacteroides* decreased after ED intervention. This phenomenon was also observed after fecal bacteria transplantation. Correspondingly, microbial translocation towards intestinal lymph nodes was inhibited in both recipient and donor mice. These results strongly supported our view that ED can reduce the abundance of mucolytic bacteria and prevent harmful microorganisms from invading intestinal epithelial cells.

In addition to mucolytic bacteria, we also found changes in the abundance of other bacteria. ED treatment increased the relative abundance of *Muribaculaceae* and *Faecalibaculum* in mice feces and decreased the relative abundance of *Alistipes*. In terms of pathogenicity, there is evidence that *Alistipes*, a newer subclass of *Bacteroidetes*, is positively correlated with diarrhea and abdominal pain, and is pathogenic in colorectal cancer (35). Therefore, the decrease in the relative abundance of intestinal *Alistipes* in mice fed with ED is related to the reduction of susceptibility to colitis. *Muribaculaceae* plays multiple functions in degrading complex carbohydrates (39). Similar to *Bifidobacterium* and *Lactobacillus*, *Faecalibaculum* can produce lactic acid and short-chain fatty acids and has anti-colon cancer effects (40). It protects the stability of the intestinal environment and prevents pathogens from colonizing in the intestinal epithelium, which has health benefits. Therefore, the higher abundance of *Muribaculaceae*, and *Faecalibaculum* and lower abundance of *Alistipes* in the intestine of mice fed with ED may also help reduce the pathogenesis of colitis. Furthermore, we observed decreased content of H_2_S and ammonia in the feces of mice treated with ED. The result suggests that the modulation of gut microbiota is likely to result from a reduction in the fermentation of unabsorbed proteins.

Endogenous endotoxins are produced by gut microbes, and enter the blood circulation with the disruption of the gut barrier (41). The LPS-LBP complex binds to TLR4 and activates MAPK and NF-κB signaling pathways (42). Previous studies have shown that colitis can be alleviated by inhibiting NF-κB or MAPK signaling pathway and reducing the release of downstream pro-inflammatory cytokines (43, 44). In this study, we found that ED significantly reduced the serum LPS level in colitis mice. The phosphorylation levels of NF-kB P65, P38, and ERK were significantly down-regulated, indicating that ED exerts an anti-inflammatory effect by preventing LPS from passing the intestinal epithelial barrier, and inhibiting NF-κB / MAPK inflammatory signal pathway. Moreover, NLRP3 inflammasome has been found to activate the expression of IL-1β and IL-18, thereby promoting the progression of IBD (45). IL-12 activates Th1 differentiation and IFN-γ release to promote intestinal mucosal inflammation (46). IL-6 and IL-23 can stimulate Th17 cells to produce IL-17 family cytokines (47), which are important regulators of intestinal mucosal inflammation. In this study, ED decreased the expression of IL-1β, IL-12, and IFN-γ. The expression of IL-12 and IL-23 were also down-regulated, but there was no difference in the expression of IL-17. The results indicate that ED may exert the protective effect through the inhibition of NLRP3 inflammasome activation and Th1 differentiation, rather than Th17 differentiation.

## Conclusion

In conclusion, ED has better preventive effects on IBD than polymeric diets. By utilizing fecal microbiota transplantation and pseudo-sterile animals, we convincingly demonstrated that gut microbiota plays a critical role in the effects of ED. ED can reduce the abundance of mucus-degrading bacteria, thereby inhibiting the mucus layer disruption and preventing harmful microbes from invading intestinal epithelium. This study may provide new insights into the gut microbiota-diet interaction in human health.

## Materials and Methods

### Reagents

Dextran sulfate sodium (DSS, MW 36-50 kDa) was obtained from MP Biomedicals LLC (Santa Ana), TRIzol^TM^ Reagent (Thermo Fisher Scientific), LunaScript^TM^SuperMix Kit (New England BioLabs), SYBR qPCR Master Mix (ChamQ^TM^Universal) were used for RT-qPCR analysis. Antibody (β-actin, p38, p-p38, JNK, p-JNK, Erk, p-Erk, p-p65, p65, occludin, ZO-1, Claudin-1, MUC-2, caspase-3, andγ-H2AX) was bought from Cell Signaling Technology; MPO kit and ELISA kits of LPS was bought from Nanjing Jiancheng Bioengineering Institute (Nanjing Jiancheng Bioengineering Institute)

### Animals

Specific-pathogen-free (SPF) ICR mice (male, 6 weeks old) were purchased from Beijing Charles River Laboratory Animal Technology Co. Ltd. They were placed in an environment with controlled temperature (25 ± 1°C) and relative humidity (50 ± 60%) with free access to food and water. The adaptation period is one week. All animal experimental protocols were approved by the Institutional Animal Care and Use Committee of Nankai University and carried out following the national ethical guidelines for laboratory animals (permission number: SYKX 2019-0001).

### Animal experimental design and Diet

One week after acclimation, the mice were weighed and randomly divided into four groups: Casein (CA) diet group, amino acid-based (AA) ED group, CA-DSS group, and AA-DSS group, with 12 mice in each group. The CA group mice were fed an AIN-93G standard diet. In the AA diet, the casein in AIN-93G was replaced with amino acids of the same composition. All diets were adjusted to the same energy level. The diet composition was listed in Table 1, and the amino acid profile of casein was shown in Table S1.

**Table 1.** Composition of experimental diets.

| Ingredient (g/kg diet) | CA | AA |
| --- | --- | --- |
| Corn Starch | 397.5 | 399.1 |
| Maltodextrin | 132 | 132 |
| Cellulose | 50 | 50 |
| Soybean Oil | 70 | 70 |
| Mineral Mix | 3.5 | 3.5 |
| Sodium Bicarbonate | 0 | 7.5 |
| Vitamin Mix | 10 | 10 |
| Choline Bitartrate | 2.5 | 2.5 |
| L-Cystine | 3 | 0 |
| Amino acid Mix | 0 | 192 |
| Casein | 200 | 0 |

After feeding all mice a specific diet for 2 weeks, two groups were treated with DSS (1.5%) in drinking water for 3 cycles (5 days/cycle, with a 7 days recovery after each of the first 2 cycles). Food consumption, body weight, and DAI were recorded regularly. At the end of the third round of DSS, mouse feces were collected, and the mice were euthanized to obtain serum. Colon tissues were immediately collected, weighed, photographed, and stored at -80°C.

### ED supplementation for antibiotic-treated mice

To deplete the intestinal flora, mice were treated with water containing a mixture of antibiotics (metronidazole 1 g/L, neomycin 1 g/L, vancomycin 500 mg/mL, ampicillin 1 g/L) for 14 days. To observe bacterial depletion, fecal DNA was isolated, and measured for the universal bacterial 16S rRNA gene by real-time qPCR. Additionally, mice were treated with antibiotics and 2.5% DSS for 7 days and then sacrificed. Mice were divided into two groups, fed with casein diet (ABX-CA) and ED (ABX-AA).

### Fecal microbiota transplantation

The donor mice were fed with a casein diet (CA) or ED (AA) for 2 weeks. Fresh feces from CA or AA were collected and resuspended in PBS (10% w/v). Fecal homogenates were centrifuged at 800×g for 5 minutes and the supernatant was collected. Fecal supernatants were orally gavaged into germ-free (GF) mice. To deplete the microbiota, the recipient mice were treated with an antibiotic cocktail (metronidazole 1g/L, neomycin 1 g/L, vancomycin 500 mg/ mL, ampicillin 1 g/L) for 2 weeks. Depletion of microbiota was confirmed by bacterial colony assays and real-time PCR analysis of universal bacterial 16S rRNA as described above. Two weeks after fecal transplantation, mice were administered with 2.5% DSS to induce colitis.

### Disease Activity Index

Disease activity index (DAI) scores were conducted periodically to assess the severity of colitis. Scoring indicators include: (a) body weight loss, (b) stool consistency, and (c) gross bleeding. Each indicator was scored as follows: change in body weight loss( 0, none; 1, 1−5%; 2, 5−10%; 3, 10−20%; 4, > 20%), stool blood (0, negative; 1, +; 2, ++; 3, +++; 4, ++++) and stool consistency (0, normal; 1 and 2, loose stool; 3 and 4, diarrhea) (48).

### Histopathological Analysis

The distal colon tissue (5 mm) was fixed in 10% formalin solution for 24 hours, then embedded in paraffin, sectioned, and stained with hematoxylin and eosin (H&E). Blind photographs were taken of each colon sample under a microscope at 100× magnification. Histological scores were examined according to the severity of crypt depletion and distortion (0–3), the degree of inflammatory infiltration (0–4), and the area of involvement (0–4) (49).

### Real-time polymerase chain reaction (RT-PCR)

Total RNA was extracted from the tissues (50-100 mg) of feces (50 mg) using TRIzol^TM^ Reagent (Thermo Fisher Scientific). The total RNA was quantified by nanodrop and then reversed to cDNA. Real-time PCR was measured using the SYBR green master mix (NEB). Cq was used to calculate the relative expression of the target gene by the 2^-ΔΔ^ Cq method. Primers for mRNA expression measurement were listed in Table S2, and primers for the measurement of bacterial 16S rRNA genes were listed in Table S3.

### Western blot analysis and Enzyme-Linked Immunosorbent Analysis (ELISA)

Colon tissue was triturated with an inhibitor cocktail (Beyotime, Beijing, China, 1:50, v/v) in RIPA lysis buffer to obtain a protein solution. After centrifugation, the protein concentration was determined by a BCA kit. The samples were separated on 12% SDS-PAGE, then transferred to the nitrocellulose filter (NC) membrane and incubated with antibodies. Membranes were washed three times with TBST and developed using the Pierce ECL western blotting kit (32106, Thermo Fisher). Antibodies were diluted at 1:1000. Secondary antibodies were used horseradish peroxidase (HRP)-conjugated secondary antibodies (1:4000 dilution, 31460, Thermo Fisher) for 1 h at room temperature.

The level of lipopolysaccharide (LPS) in serum was detected with an ELISA kit. The activity of MPO in colon tissue was determined with a commercially available kit.

### Immunofluorescence Analysis

The paraffin-embedded samples were sectioned. The prepared sections are fluorescently stained in the order of section dewaxing, protease K treatment, denaturation, hybridization, sealing, DAPI re-staining, and anti-fluorescent quencher sealing. The experimental procedure was protected from light, and then a microscopic examination was carried out. Three fields of view were randomly selected for each colon sample under 100× and 200× magnification.

### 16S rRNA high-throughput sequencing and bioinformatics analysis

High-throughput sequencing and bioinformatics analysis were performed according to our previous method (50), and were presented in the supplementary material. The data were deposited in the NCBI Sequence Read Archive, with an accession number of PRJNA841685.

### Determination of Mucus Degrading Enzyme Activity

The activity of bacteria-derived mucus-degrading enzymes and carbohydrate-active enzymes were determined according to the previously described method with minor modifications (51). Approximately 50 mg of feces were homogenized in enzyme analysis buffer (50 mM Tris, 100 mM KCl, 10 mM MgCl_2_, pH 7.26). 100 uL of 12% Triton X-100 buffer, lysozyme, and 10 mg of DNase and protease inhibitor (Roche) were added to the mixture and homogenized on ice to obtain a fecal suspension. The suspension was centrifuged at 10,000×g for 10 minutes, and the obtained supernatant was used for enzyme activity assay. 10 μL sample was mixed with 150 μL of 10 nM substrates solution, including 4-nitrophenyl-2-acetamino-2-deoxy-β-D-glucopyranoside, 4-nitrophenylsulfate, 4-nitrophenyl-α-L-fucopyranoside, 4-nitrophenyl-α-D-galactose-pyran-side, 4-nitrophenyl-β-D-glucopyranoside, and 4-nitrophenyl-β-D-xylopyranoside prepared in the same assay buffer. The mixtures were incubated at 37℃ for 2 h, and the absorbance at 405 nm was recorded. A standard curve was established with 4-nitrophenol.

### Assessment of bacterial translocation in mesenteric lymph nodes

Microbial translocation in the mesenteric lymph nodes was detected according to the method previously described (52). Under anaerobic conditions, MLN were collected aseptically and homogenized in BBL Mycoflask Thioglycollate (liquid) Prepared Media (BD Diagnostic Systems). Gifu medium agar plates containing the contents of the BBL Mycoflask Thioglycollate were incubated in an anaerobic incubator at 37°C for 48 h. Colony formation units were counted, and the concentration per milligram of tissue was determined.

### Determination of fecal content of H_2_S and ammonia

To determine the fecal content of H_2_S and ammonia, 50 mg of feces were homogenized in 500 μL of 0.1 M HCl. The samples were centrifuged at 12,000×g for 20 min. The concentration of H_2_S was determined by the N, N-dimethyl-p-phenylenediamine sulfate method (53). The concentration of ammonia was determined by the phenol-hypochlorite method (54).

### Statistical Analysis

Statistical analysis was performed using GraphPad Prism 8.0. Data were presented as the mean ± SEM. Differences among groups were compared by one-way ANOVA followed by Bonferroni post hoc test. Differences between two groups were analyzed by two-tailed unpaired Student’s t-test or Mann-Whitney test. *P*<0.05 was considered statistically significant.

## Acknowledgments

This work was funded by the National Natural Science Foundation of China (32101962).

Bowei Zhang: Investigation, Data analysis, Writing-original draft, Funding acquisition. Congying Zhao: Investigation, Data analysis, Writing-original draft. Yunhui Zhang: Investigation. Xuejiao Zhang: Investigation. Xiang Li: Methodology. Xiaoxia Liu: Investigation. Jia Yin: Methodology. Xinyang Li: review. Jin Wang: review. Shuo Wang: Supervision, Project administration, Writing-review & editing.

The authors declare no competing interests.

## Supplementary material

Supplementary Materials and Methods. Table S1, amino acid profile of casein. Table S2, primers for mRNA expression measurement in RT-qPCR. Table S3, primers for the measurement of bacterial 16S rRNA genes by RT-qPCR. Figure S1, statistical analysis of PCoA. Figure S2, difference between gut microbiota in the CA group and the AA group. Figure S3, effect of antibiotics cocktail on depleting the gut microbiota in mice. Figure S4, relative mRNA expression of gut barrier-related proteins in antibiotic-treated pseudo-germ-free mice.

## Reference

1. Ananthakrishnan AN. 2013. Environmental risk factors for inflammatory bowel disease. Gastroenterology & hepatology 9:367–374.

2. Kaplan GG. 2015. The global burden of IBD: from 2015 to 2025. Nat Rev Gastroenterol Hepatol 12:720–727.

3. Kaplan GG, Ng SC. 2017. Understanding and Preventing the Global Increase of Inflammatory Bowel Disease. Gastroenterology 152:313-+.

4. Zhao B, Xia B, Li X, Zhang L, Liu X, Shi R, Kou R, Liu Z, Liu X. 2020. Sesamol Supplementation Attenuates DSS-Induced Colitis via Mediating Gut Barrier Integrity, Inflammatory Responses, and Reshaping Gut Microbiome. J Agric Food Chem 68:10697–10708.

5. Ryan FJ, Ahern AM, Fitzgerald RS, Laserna-Mendieta EJ, Power EM, Clooney AG, O’Donoghue KW, McMurdie PJ, Iwai S, Crits-Christoph A, Sheehan D, Moran C, Flemer B, Zomer AL, Fanning A, O’Callaghan J, Walton J, Temko A, Stack W, Jackson L, Joyce SA, Melgar S, DeSantis TZ, Bell JT, Shanahan F, Claesson MJ. 2020. Colonic microbiota is associated with inflammation and host epigenomic alterations in inflammatory bowel disease. Nature Communications 11.

6. Desai MS, Seekatz AM, Koropatkin NM, Kamada N, Hickey CA, Wolter M, Pudlo NA, Kitamoto S, Terrapon N, Muller A, Young VB, Henrissat B, Wilmes P, Stappenbeck TS, Nunez G, Martens EC. 2016. A Dietary Fiber-Deprived Gut Microbiota Degrades the Colonic Mucus Barrier and Enhances Pathogen Susceptibility. Cell 167:1339–1353 e1321.

7. Johansson MEV, Hansson GC. 2016. Immunological aspects of intestinal mucus and mucins. Nature Reviews Immunology 16:639–649.

8. Hansson GC, Johansson MEV. 2010. The inner of the two Muc2 mucin-dependent mucus layers in colon is devoid of bacteria. Gut Microbes 1:51–54.

9. Paone P, Cani PD. 2020. Mucus barrier, mucins and gut microbiota: the expected slimy partners? Gut 69:2232–2243.

10. Johansson MEV, Gustafsson JK, Holmen-Larsson J, Jabbar KS, Xia L, Xu H, Ghishan FK, Carvalho FA, Gewirtz AT, Sjovall H, Hansson GC. 2014. Bacteria penetrate the normally impenetrable inner colon mucus layer in both murine colitis models and patients with ulcerative colitis. Gut 63:281–291.

11. Png CW, Linden SK, Gilshenan KS, Zoetendal EG, McSweeney CS, Sly LI, McGuckin MA, Florin THJ. 2010. Mucolytic Bacteria With Increased Prevalence in IBD Mucosa Augment In Vitro Utilization of Mucin by Other Bacteria. American Journal of Gastroenterology 105:2420–2428.

12. Luis AS, Jin C, Pereira GV, Glowacki RWP, Gugel SR, Singh S, Byrne DP, Pudlo NA, London JA, Basle A, Reihill M, Oscarson S, Eyers PA, Czjzek M, Michel G, Barbeyron T, Yates EA, Hansson GC, Karlsson NG, Cartmell A, Martens EC. 2021. A single sulfatase is required to access colonic mucin by a gut bacterium. Nature 598:332-+.

13. Ndeh D, Basle A, Strahl H, Yates EA, McClurgg UL, Henrissat B, Terrapon N, Cartmell A. 2020. Metabolism of multiple glycosaminoglycans by Bacteroides thetaiotaomicron is orchestrated by a versatile core genetic locus. Nat Commun 11:646.

14. Johansson MEV, Sjovall H, Hansson GC. 2013. The gastrointestinal mucus system in health and disease. Nature Reviews Gastroenterology & Hepatology 10:352–361.

15. Adam L, Phulukdaree A, Soma P. 2018. Effective long-term solution to therapeutic remission in Inflammatory Bowel Disease: Role of Azathioprine. Biomed Pharmacother 100:8–14.

16. Arebi N, Dyall L, Kamperidis N. 2021. A User’s Guide to De-Escalating Immunomodulator and Biologic Therapy in Inflammatory Bowel Disease. Clin Gastroenterol Hepatol 19:1300–1301.

17. Dolan KT, Chang EB. 2017. Diet, gut microbes, and the pathogenesis of inflammatory bowel diseases. Mol Nutr Food Res 61.

18. Sasson AN, Ananthakrishnan AN, Raman M. 2021. Diet in Treatment of Inflammatory Bowel Diseases. Clin. Gastroenterol. Hepatol. 19:425–435 e423.

19. Lan A, Blachier F, Benamouzig R, Beaumont M, Barrat C, Coelho D, Lancha A, Jr., Kong X, Yin Y, Marie JC, Tome D. 2015. Mucosal healing in inflammatory bowel diseases: is there a place for nutritional supplementation? Inflamm. Bowel Dis. 21:198–207.

20. Sugihara K, Morhardt TL, Kamada N. 2018. The Role of Dietary Nutrients in Inflammatory Bowel Disease. Front. Immunol. 9:3183.

21. Menezes-Garcia Z, Kumar A, Zhu W, Winter SE, Sperandio V. 2020. l-Arginine sensing regulates virulence gene expression and disease progression in enteric pathogens. Proc. Natl. Acad. Sci. U. S. A. 117:12387–12393.

22. Lee D, Albenberg L, Compher C, Baldassano R, Piccoli D, Lewis JD, Wu GD. 2015. Diet in the pathogenesis and treatment of inflammatory bowel diseases. Gastroenterology 148:1087–1106.

23. Andou A, Hisamatsu T, Okamoto S, Chinen H, Kamada N, Kobayashi T, Hashimoto M, Okutsu T, Shimbo K, Takeda T, Matsumoto H, Sato A, Ohtsu H, Suzuki M, Hibi T. 2009. Dietary histidine ameliorates murine colitis by inhibition of proinflammatory cytokine production from macrophages. Gastroenterology 136:564–574 e562.

24. Guo D, Yang J, Ling F, Tu L, Li J, Chen Y, Zou K, Zhu L, Hou X. 2020. Elemental Diet Enriched with Amino Acids Alleviates Mucosal Inflammatory Response and Prevents Colonic Epithelial Barrier Dysfunction in Mice with DSS-Induced Chronic Colitis. Journal of immunology research 2020:9430763.

25. Giaffer MH, North G, Holdsworth CD. 1990. Controlled trial of polymeric versus elemental diet in treatment of active Crohn’s disease. Lancet 335:816–819.

26. Rigaud D, Cosnes J, Le Quintrec Y, Rene E, Gendre JP, Mignon M. 1991. Controlled trial comparing two types of enteral nutrition in treatment of active Crohn’s disease: elemental versus polymeric diet. Gut 32:1492–1497.

27. Lavelle A, Sokol H. 2020. Gut microbiota-derived metabolites as key actors in inflammatory bowel disease. Nature Reviews Gastroenterology & Hepatology 17:223–237.

28. Wastyk HC, Fragiadakis GK, Perelman D, Dahan D, Merrill BD, Yu FB, Topf M, Gonzalez CG, Van Treuren W, Han S, Robinson JL, Elias JE, Sonnenburg ED, Gardner CD, Sonnenburg JL. 2021. Gut-microbiota-targeted diets modulate human immune status. Cell 184:4137–4153 e4114.

29. Johansson ME, Gustafsson JK, Holmen-Larsson J, Jabbar KS, Xia L, Xu H, Ghishan FK, Carvalho FA, Gewirtz AT, Sjovall H, Hansson GC. 2014. Bacteria penetrate the normally impenetrable inner colon mucus layer in both murine colitis models and patients with ulcerative colitis. Gut 63:281–291.

30. Van der Sluis M, De Koning BA, De Bruijn AC, Velcich A, Meijerink JP, Van Goudoever JB, Buller HA, Dekker J, Van Seuningen I, Renes IB, Einerhand AW. 2006. Muc2-deficient mice spontaneously develop colitis, indicating that MUC2 is critical for colonic protection. Gastroenterology 131:117–129.

31. Yang K, Popova NV, Yang WC, Lozonschi I, Tadesse S, Kent S, Bancroft L, Matise I, Cormier RT, Scherer SJ, Edelmann W, Lipkin M, Augenlicht L, Velcich A. 2008. Interaction of Muc2 and Apc on Wnt signaling and in intestinal tumorigenesis: Potential role of chronic inflammation. Cancer Research 68:7313–7322.

32. Yao D, Dai W, Dong M, Dai C, Wu S. 2021. MUC2 and related bacterial factors: Therapeutic targets for ulcerative colitis. EBioMedicine 74:103751.

33. Luis AS, Jin C, Pereira GV, Glowacki RWP, Gugel SR, Singh S, Byrne DP, Pudlo NA, London JA, Basle A, Reihill M, Oscarson S, Eyers PA, Czjzek M, Michel G, Barbeyron T, Yates EA, Hansson GC, Karlsson NG, Cartmell A, Martens EC. 2021. A single sulfatase is required to access colonic mucin by a gut bacterium. Nature 598:332–337.

34. David LA, Maurice CF, Carmody RN, Gootenberg DB, Button JE, Wolfe BE, Ling AV, Devlin AS, Varma Y, Fischbach MA, Biddinger SB, Dutton RJ, Turnbaugh PJ. 2014. Diet rapidly and reproducibly alters the human gut microbiome. Nature 505:559-+.

35. Saulnier DM, Riehle K, Mistretta T-A, Diaz M-A, Mandal D, Raza S, Weidler EM, Qin X, Coarfa C, Milosavljevic A, Petrosino JF, Highlander S, Gibbs R, Lynch SV, Shulman RJ, Versalovic J. 2011. Gastrointestinal Microbiome Signatures of Pediatric Patients With Irritable Bowel Syndrome. Gastroenterology 141:1782–1791.

36. Yoon HS, Cho CH, Yun MS, Jang SJ, You HJ, Kim J-h, Han D, Cha KH, Moon SH, Lee K, Kim Y-J, Lee S-J, Nam T-W, Ko G. 2021. Akkermansia muciniphila secretes a glucagon-like peptide-1-inducing protein that improves glucose homeostasis and ameliorates metabolic disease in mice. Nature Microbiology 6:563-+.

37. Seregin SS, Golovchenko N, Schaf B, Chen J, Pudlo NA, Mitchell J, Baxter NT, Zhao L, Schloss PD, Martens EC, Eaton KA, Chen GY. 2017. NLRP6 Protects Il10(-/-) Mice from Colitis by Limiting Colonization of Akkermansia muciniphila. Cell Rep 19:2174.

38. Png CW, Lindén SK, Gilshenan KS, Zoetendal EG, McSweeney CS, Sly LI, McGuckin MA, Florin THJ. 2010. Mucolytic Bacteria With Increased Prevalence in IBD Mucosa AugmentIn VitroUtilization of Mucin by Other Bacteria. Official journal of the American College of Gastroenterology | ACG 105:2420–2428.

39. Lagkouvardos I, Lesker TR, Hitch TCA, Galvez EJC, Smit N, Neuhaus K, Wang J, Baines JF, Abt B, Stecher B, Overmann J, Strowig T, Clavel T. 2019. Sequence and cultivation study of Muribaculaceae reveals novel species, host preference, and functional potential of this yet undescribed family. Microbiome 7:28.

40. Fuhren J, Schwalbe M, Boekhorst J, Rosch C, Schols HA, Kleerebezem M. 2021. Dietary calcium phosphate strongly impacts gut microbiome changes elicited by inulin and galacto-oligosaccharides consumption. Microbiome 9:218.

41. Stephens M, von der Weid PY. 2020. Lipopolysaccharides modulate intestinal epithelial permeability and inflammation in a species-specific manner. Gut Microbes 11:421–432.

42. Kayagaki N, Wong MT, Stowe IB, Ramani SR, Gonzalez LC, Akashi-Takamura S, Miyake K, Zhang J, Lee WP, Muszynski A, Forsberg LS, Carlson RW, Dixit VM. 2013. Noncanonical Inflammasome Activation by Intracellular LPS Independent of TLR4. Science 341:1246–1249.

43. Papoutsopoulou S, Campbell BJ. 2021. Epigenetic Modifications of the Nuclear Factor Kappa B Signalling Pathway and its Impact on Inflammatory Bowel Disease. Current Pharmaceutical Design 27:3702–3713.

44. Li S, Ma B, Wang J, Peng H, Zheng M, Dai W, Liu J. 2020. Novel Pentapeptide Derived from Chicken by-Product Ameliorates DSS-Induced Colitis by Enhancing Intestinal Barrier Function via AhR-Induced Src Inactivation. J Agric Food Chem.

45. Zhen Y, Zhang H. 2019. NLRP3 Inflammasome and Inflammatory Bowel Disease. Front. Immunol. 10:276.

46. Moschen AR, Tilg H, Raine T. 2019. IL-12, IL-23 and IL-17 in IBD: immunobiology and therapeutic targeting. Nat Rev Gastroenterol Hepatol 16:185–196.

47. Gunasekera DC, Ma J, Vacharathit V, Shah P, Ramakrishnan A, Uprety P, Shen Z, Sheh A, Brayton CF, Whary MT, Fox JG, Bream JH. 2020. The development of colitis in Il10(-/-) mice is dependent on IL-22. Mucosal Immunol 13:493–506.

48. Han Y, Song M, Gu M, Ren D, Zhu X, Cao X, Li F, Wang W, Cai X, Yuan B, Goulette T, Zhang G, Xiao H. 2019. Dietary Intake of Whole Strawberry Inhibited Colonic Inflammation in Dextran-Sulfate-Sodium-Treated Mice via Restoring Immune Homeostasis and Alleviating Gut Microbiota Dysbiosis. J Agric Food Chem 67:9168–9177.

49. Zhang B, Xu Y, Liu S, Lv H, Hu Y, Wang Y, Li Z, Wang J, Ji X, Ma H, Wang X, Wang S. 2020. Dietary Supplementation of Foxtail Millet Ameliorates Colitis-Associated Colorectal Cancer in Mice via Activation of Gut Receptors and Suppression of the STAT3 Pathway. Nutrients 12.

50. Zhang B, Xu Y, Lv H, Pang W, Wang J, Ma H, Wang S. 2021. Intestinal pharmacokinetics of resveratrol and regulatory effects of resveratrol metabolites on gut barrier and gut microbiota. Food Chem 357:129532.

51. Khan S, Waliullah S, Godfrey V, Khan MAW, Ramachandran RA, Cantarel BL, Behrendt C, Peng L, Hooper LV, Zaki H. 2020. Dietary simple sugars alter microbial ecology in the gut and promote colitis in mice. Sci Transl Med 12.

52. Manfredo Vieira S, Hiltensperger M, Kumar V, Zegarra-Ruiz D, Dehner C, Khan N, Costa FRC, Tiniakou E, Greiling T, Ruff W, Barbieri A, Kriegel C, Mehta SS, Knight JR, Jain D, Goodman AL, Kriegel MA. 2018. Translocation of a gut pathobiont drives autoimmunity in mice and humans. Science 359:1156–1161.

53. Cline JD. 1969. SPECTROPHOTOMETRIC DETERMINATION OF HYDROGEN SULFIDE IN NATURAL WATERS1. Limnology & Oceanography 14.

54. Solórzano L. 1969. DETERMINATION OF AMMONIA IN NATURAL WATERS BY THE PHENOLHYPOCHLORITE METHOD 1 1 This research was fully supported by U.S. Atomic Energy Commission Contract No. ATS (11-1) GEN 10, P.A. 20. Limnology & Oceanography 14:799–801.

